# Intracellular passage of Na^+^ in an active state g-protein coupled receptor

**DOI:** 10.1101/124420

**Authors:** Owen N. Vickery, Catarina A. Carvalheda, Saheem A. Zaidi, Andrei V. Pisliakov, Vsevolod Katritch, Ulrich Zachariae

## Abstract

Playing a central role in cell signalling, GPCRs have evolved into the largest superfamily of membrane proteins and form the majority of drug targets in humans. How extracellular agonist binding triggers the activation of GPCRs and associated intracellular effector proteins remains, however, poorly understood. High resolution structural studies have recently revealed that inactive class-A GPCRs harbour a conserved binding site for Na^+^ ions in the centre of their transmembrane domain, accessible from the extracellular space. Here, we show that the opening of a conserved hydrated channel in the activated state receptors allows the Na^+^ ion to egress from its binding site into the cytosol. Coupled with protonation changes, this ion movement occurs without significant energy barriers, and can be driven by physiological transmembrane ion and voltage gradients. We propose that Na^+^ ion exchange with the cytosol is a key step in GPCR activation, which locks receptors in long-lived active-state conformations.

## INTRODUCTION

G-protein coupled receptors (GPCRs) mediate the transfer of external ligand binding information across the plasma membrane to activate a range of intracellular signaling pathways (Pierce, Premont, & Lefkowitz, 2002). Playing a central role in regulation of vital biological systems, including nervous, cardiovascular, immune, digestive, reproductive etc., they represent the majority of membrane proteins in humans and the largest class of present drug targets (Overington, Al-Lazikani, & Hopkins, 2006; Rask-Andersen, Masuram, & Schiöth, 2014). In recent years, a number of crystal structures have been solved to reveal conformational changes between inactive and active state receptors, including common movement in transmembrane helices and conserved microswitches (Katritch, Cherezov, & Stevens, 2013; Venkatakrishnan et al., 2013). However, despite this wealth of structural information, it is still not fully understood how ligand binding leads to activated receptors, which are able to trigger nucleotide exchange in intracellular effector G-protein complexes.

One of the major unknowns is the role of the highly conserved hydrophilic water-filled channel observed in crystal structures of class A GPCRs, which extends along the receptor axis from the external ligand binding region nearly all the way to the effector binding site. The channel is sealed towards the cytoplasm by a thin layer of hydrophobic residues in inactive state GPCRs (Fig 1A,B). Structures of high resolution, crystallized in the inactive conformation, reveal a Na^+^ ion near the floor of this pocket, coordinated by water and three or four conserved residues including an acidic aspartate that is fully conserved in all ligand-sensing class A GPCRs (Christopher et al., 2013; Fenalti et al., 2014; Kruse et al., 2012; Liu et al., 2012; Miller-Gallacher et al., 2014; Pardo, Deupi, Dölker, López-Rodríguez, & Campillo, 2007; Zhang et al., 2012) (D^2.50^; superscript refers to the Ballesteros and Weinstein residue numbering system) (Isberg et al., 2015). The allosteric effect of monovalent cations, in particular Na^+^ ions, for GPCR function has been known for almost half a century (Pert & Synder, 1974), and the bulk of recent evidence shows that these effects are largely mediated by the ion binding at the D^2.50^ site at the physiological concentration of Na^+^ (140 mM and lower) (Fenalti et al., 2014; Liu et al., 2012; Massink et al., 2015). Due to the highly conserved nature of D^2.50^ and other Na^+^ ion coordinating residues, Na^+^ ion binding at this site is likely to be a ubiquitous feature shared by the vast majority of class A GPCRs (Katritch et al., 2014).

**Figure 1:**
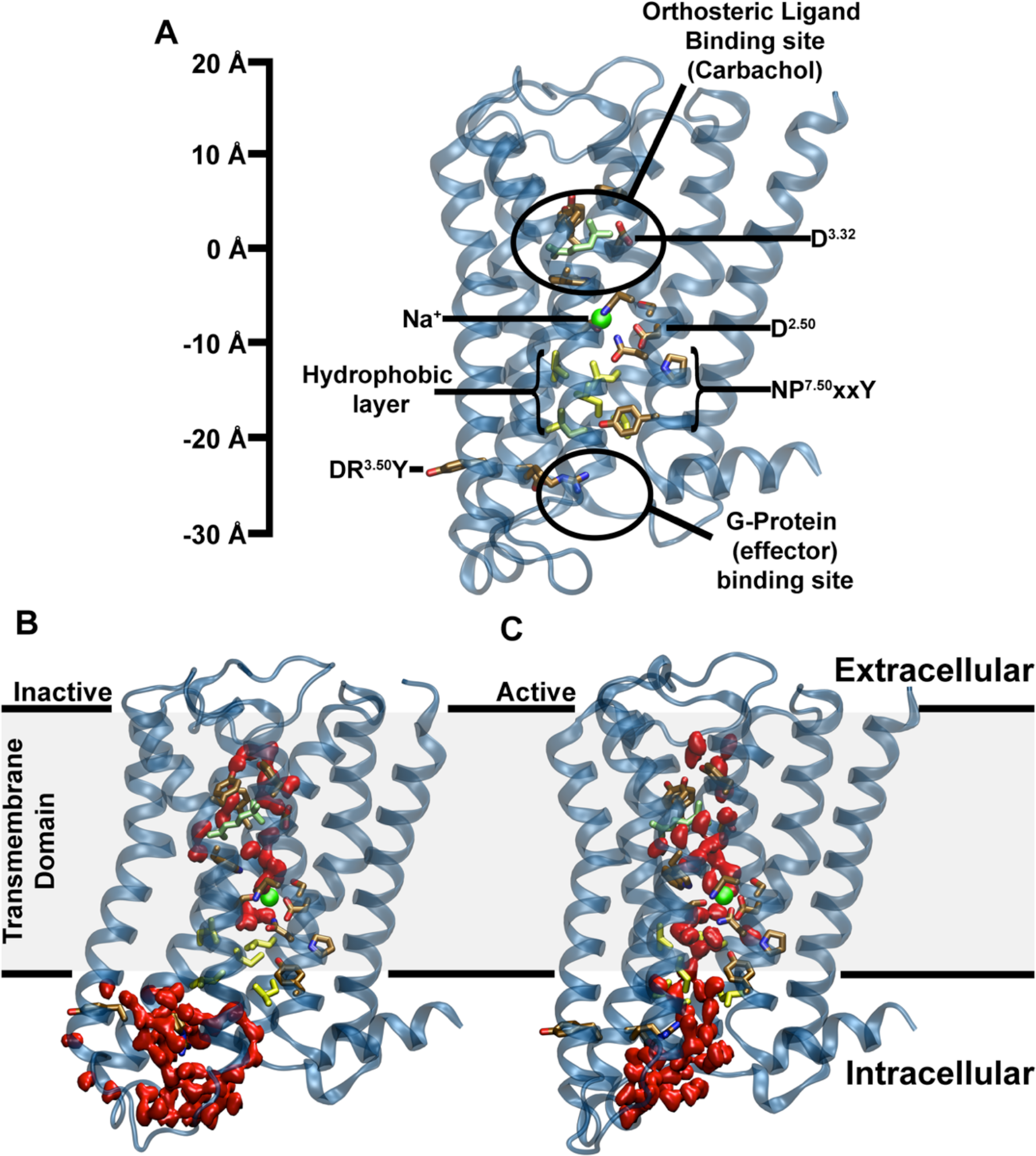
Major structural features and internal hydration of class A GPCRs in the inactive and active state as shown by the m2r. **(A)** The main structural features of class A GPCRs, as exemplified by the m2r, include 7 TM helices (blue), an extracellular ligand binding site, the intracellular effector (G-protein) binding site as well as conserved and functionally important residues termed microswitches (selected ones are highlighted). The scale bar shown and all positions stated in the text use the Cα atom of D103^3.32^ as reference. **(B)** Conformation of inactive m2r (PDB: 3UON) during the simulations showing the presence of the hydrophobic layer separating the hydrophilic pocket and effector binding site. **(C)** After transition to the active state (PDB: 4MQT), and further simulation, m2r displays a continuous water channel connecting the orthosteric ligand binding site, hydrophilic pocket and effector binding site. Water molecules are shown in red (surface representation), the position of the allosteric Na^+^ ion, as obtained from our initial simulations, is shown as a green sphere, and residues forming the hydrophobic layer (yellow) as well as the bound ligand (carbachol, light green) are depicted in stick representation.

In active receptor conformations, the ion binding site near D^2.50^ shows a collapsed state, which is likely not optimal for Na^+^ ion binding (Huang et al., 2015; Kruse et al., 2013; Liu et al., 2012; Rasmussen et al., 2011). It was therefore proposed that Na^+^ ion leaves the hydrophilic pocket upon receptor activation by a ligand or during receptor-G-protein complex formation. However, how this movement is triggered and which pathway is followed by the ion remains unknown.

Here, we investigated the link between ligand-induced receptor activation, the fate of the bound Na^+^ ion in class A GPCRs and its implications for transmembrane (TM) signal transduction by equilibrium and non-equilibrium atomistic simulations on the M2 muscarinic receptor (m2r). When one addresses these questions, it is important to take physiologically relevant electrochemical membrane conditions into consideration. Strong TM Na^+^ and K^+^ gradients produce a sizable voltage across the plasma membrane of up to -100 mV in the resting state of mammalian cells (Kandel, Schwartz, & Jessell, 2000). Both the ionic gradients and electric field have been shown to influence the function of GPCRs (Ben-Chaim et al., 2006; Navarro-Polanco et al., 2011; Rinne, Mobarec, Mahaut-Smith, Kolb, & Bunemann, 2015) and are likely to impact the movement of the Na^+^ ion within the membrane region.

Our data reveal that the Na^+^ ion observed in the TM domain of class A GPCRs can readily traverse the receptor and, driven by the electrochemical gradients, migrate into the cytoplasm in active receptor conformations. This result implies that a Na^+^ ion may be exchanged from the extracellular space to the cytoplasm as an important step in receptor activation. Furthermore, the movement of Na^+^ in the receptor, and intracellular egress, are coupled to a protonation change of D^2.50^.

## RESULTS

### GPCR activation opens a hydrated pathway across the receptor

We were first interested whether the conformational change from the inactive to active receptor state renders the ion binding pocket sterically incapable of accommodating a Na^+^ ion. The binding site for Na^+^ appears to adopt a collapsed conformation in active crystal structures. We started from an inactive state structure of the m2 muscarinic acetylcholine receptor (m2r, PDB ID: 3UON) and, using a targeted molecular dynamics (MD) approach, gently drove this conformation to the active state of this receptor (PDB ID: 4MQT) (Fig S1).

Our simulations show that the active state of m2r initially retains sufficient space for the ion. The electrostatic attraction between the ion and the negatively charged side chain of D69^2.50^ keeps the ion bound to this site during and after the transition from the inactive to the active receptor conformation (Fig S2). However, our simulations show a widening of the intracellular portion of the TM helices below the hydrophilic pocket during this conformational change, which subsequently becomes fully hydrated (Fig 1B). The hydrated pathway forms a connection between the orthosteric ligand-binding site, the hydrophilic pocket and the G-protein binding site. The slim hydrophobic layer that delimits the hydrophilic pocket towards the G-protein binding site in the inactive crystal structure undergoes substantial conformational changes, which are especially evident from the sidechain position of Y440^7.53^. Our simulations show two major conformations of the Y440^7.53^ sidechain following the transition – an upward state similar to the conformation observed in the inactive crystal structure (PDB: 3UON; Fig S3B) and a downward configuration, which is also seen in the active crystal structure (PDB: 4MQT; Fig S3A). The formation of a hydrated pathway connecting the receptor ligand and effector binding sites has been reported in previous simulation studies on the A_2A_R and 5-HT_1A_ receptors (Yuan, Filipek, Palczewski, & Vogel, 2014; Yuan, Peng, Palczewski, Vogel, & Filipek, 2016), however the previous reports did not take the presence of a Na^+^ ion into consideration.

### The position of the internal Na^+^ ion is coupled to protonation of D2.50

We were next interested in the interplay between the Na^+^ ion and the key conserved titratable residue D69^2.50^. A number of computational studies have explored functional implications of the protonation state of D^2.50^, in particular its role in receptor activation, Na^+^ ion binding, and interaction with the “ionic lock” motif (D^3.49^R^3.50^Y^3.51^) in several class A family GPCRs (Miao, Caliman, & McCammon, 2015; Ranganathan, Dror, & Carlsson, 2014; Vanni, Neri, Tavernelli, & Rothlisberger, 2010). Here, we focused on a potential coupling between the position of the Na^+^ ion within the receptor and protonation of D69^2.50^. We carried out p*K*_a_ calculations on D69^2.50^ using more than 800 equilibrated frames from simulations of the m2r receptor in a variety of conformations, including both the upward and downward configurations of the Y440^7.53^ side chain. Due to the formation of a hydrated pathway across the receptor from the ligand to the effector binding sites in the active state simulations, we were able to evaluate the effect of the Na^+^ ion positional changes on the D69^2.50^ p*K*_a_, where the Na^+^ ion was shifted both in the upward (towards the extracellular face) and downward direction.

Figure 2 shows that the p*K*_a_ value and, thus, the most likely protonation state of D69^2.50^ are strongly influenced by the Na^+^ ion. If the cation is within ~3-5Å of D69^2.50^, its positive charge strongly stabilises the negatively charged form of D69^2.50^, leading to a p*K*_a_ value of ~3–4. However, displacement of the Na^+^ ion to distances of 5Å and greater gives rise to a substantial p*K*_a_ shift to values between 8-12. This can be understood given the location of D69^2.50^ in the middle of the transmembrane domain, surrounded by many non-polar residues. Transient movements of the internal Na^+^ ion from its binding site, facilitated by activation-related conformational changes in the Na^+^ pocket, can therefore be sufficient to lead to protonation of D69^2.50^.

**Figure 2:**
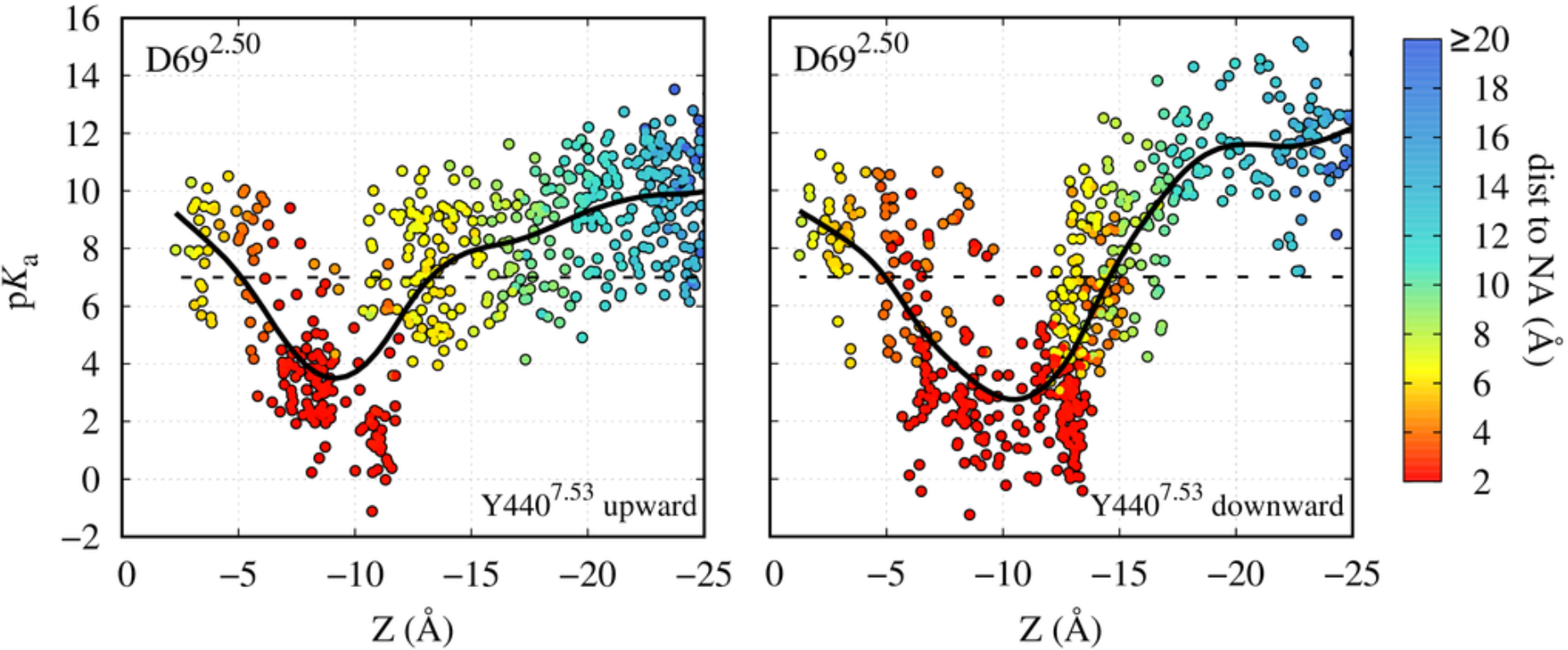
Proximity of the Na^+^ ion modulates protonation of D69^2.50^. Continuum electrostatics calculations of the p*K*_a_ of the D69^2.50^ sidechain using a multitude of m2r conformations obtained from our atomistic simulations in the carbachol-bound active state, both for Y440^7.53^ in the upward (left) and downward (right) conformations. The p*K*_a_ is shown as a function of *Z*, the separation between the Na^+^ ion and the C_α_ atom of D103^3.32^, which marks the orthosteric ligand binding pocket, along the TM axis (see Fig 1A). The data points are in addition coloured according to their distance to the D69^2.50^ sidechain. The black continuous line, a smoothed spline fit, indicates the approximate average p*K*_a_ for each separation for illustrative purposes, and the dashed black line shows a p*K*_a_ of 7.

For the protonation of D^2.50^, we propose that the most likely proton entry route would be from the extracellular side, along the negative membrane potential gradient. Moreover, in the m2r and other aminergic receptors the proton could be transferred from the conserved D^3.32^ in the orthosteric binding pocket via a short chain of water molecules (Isom & Dohlman, 2015). In the apo state, our calculations in m2r indicate that D^3.32^ is generally protonated (p*K*_a_ = 11.2±1.7), whereas upon ligand binding the p*K*_a_ is substantially lowered (p*K*_a_ = 7.6±1.9). A possible protonation change of D^3.32^ could thus facilitate the shuttling of protons to D^2.50^. Furthermore, if a G-protein complex with a receptor is preformed before agonist binding, D^2.50^ would be readily accessible for protonation from the extracellular side via a hydrated pathway. In this context, it has further been argued that bound agonists, but not antagonists, may sustain the hydrated pathway past the ligand which connects the extracellular space with the Na^+^ ion binding site upon receptor activation (Yuan et al., 2016).

### Simulations under electrochemical gradient show ion movement to the intracellular face

Next we conducted atomistic simulations with the Computational Electrophysiology (CompEL) protocol (Kutzner et al., 2016) on the active conformation of m2r. We applied a physiological Na^+^ ion gradient of 150:10mM across the membrane from the extracellular to the intracellular side, in addition to a small ion imbalance evoking a hyperpolarised V_m_ at - 250mV. Due to the wide range of p*K*_a_ values that D69^2.50^ can adopt, its sidechain was modeled both in charged and neutral forms.

Our simulations at -250 mV show that the Na^+^ ion exhibits a substantial degree of mobility even when D69^2.50^ is in the charged state (Fig 3A, B, S7). The Na^+^ ion is predominantly coordinated by the residues D69^2.50^, S110^3.39^, N435^7.45^ and S433^7.46^. Under a small membrane voltage, a bimodal distribution of distances between the ion and D69^2.50^ is observed, where larger distances of 5–6 Å are not uncommon (Fig S7C). As our p*K*_a_ calculations showed that moderate excursions of the ion from its original binding site on this scale likely have a major impact on the p*K*_a_ and protonation state of the D69^2.50^ sidechain (Fig 2), we investigated the effect of a protonation change of D69^2.50^ in the active conformation.

Our simulations reveal that, in this receptor conformation, the Na^+^ ion readily passes through the hydrated channel into the intracellular solution. When D69^2.50^ is neutral, we observe the Na^+^ ion to be expelled into the intracellular solution in three out of four simulations at -250mV (Fig 3A, C, for complete list of trajectories see S4, 7A). At -500mV the effect is, expectably, even more pronounced and movement into the cytoplasm is seen in all the simulations we conducted (Fig 3B, D, for complete list of trajectories see S4, 7A). In contrast, when D69^2.50^ is charged, such a transition is observed in only in one out of eight simulations at a raised membrane voltage (Fig 3B, S7B). The observed translocation of Na^+^ to the intracellular side occurs irrespective of the conformation adopted by Y440^7.53^ (Fig 3C, D, S3).

**Figure 3:**
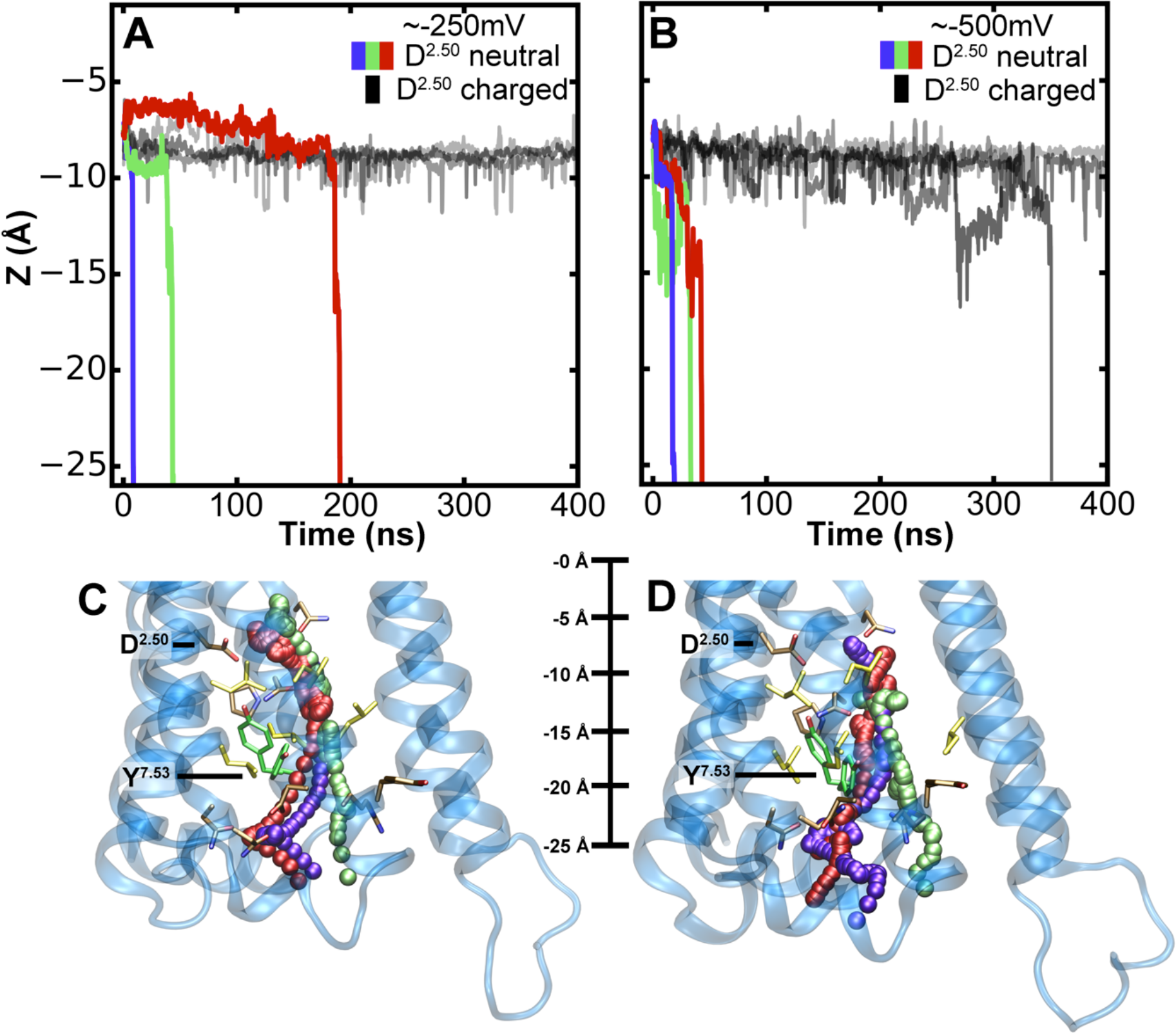
Na^+^ ion migration across the receptor to the intracellular side. Z-coordinate of the Na^+^ ion in the m2r under a hyperpolarised V_m_ of -250mV **(A)** and -500mV **(B)**. Black lines denote the simulations with a charged D69^2.50^; the purple, green and red lines display simulations with a neutral D69^2.50^. Here we show a selection of simulations, when D69^2.50^ is neutral; the trajectories show full passage of the ion to the intracellular side (for complete list of trajectories see Fig S4, S7). Trajectories of the Na^+^ ion moving from the hydrophilic pocket, accessible from the extracellular space, into the intracellular bulk solution at **(C)** -250mV and **(D)** -500mV. The colour of the Na^+^ ion corresponds to panels A and Β respectively. The Y440^7.53^ upward and downward conformations are shown in green. The pathways of the ion towards the intracellular side are almost indistinguishable from each other until the ion passes Y440^7.53^, thereafter the pathways diverge somewhat due to the widened exit region to the cytoplasm.

In our simulations as well as under physiological conditions, both TM ion concentration and voltage gradients drive ion flow across membrane pores. In the case of the Na^+^ ion, both gradients act synergistically in the resting state of the cell, driving the Na^+^ ion towards the cytoplasm. Under the conditions used in the simulations, fast ion motion through the receptor is predominantly voltage-driven. Converted into an effective force, and using a linear approximation to describe the gradient across the membrane (Dill & Bromberg, 2011), the influence of the concentration gradient would be about 10-fold smaller (~1.3 pN) than the driving force caused by the voltage gradient in these conditions (~13 pN).

### Energetics of ion movement to the cytoplasm

As the initiation of fast movement of ions to the intracellular side was initially tested under slightly supra-physiological levels of V_m_ in our CompEL simulations of active state m2r, we next evaluated the detailed equilibrium energetics of the Na^+^ ion movement on this pathway (i.e without applied gradients) to ascertain the physiological relevance of this transition. We calculated the potential-of-mean-force (PMF) for the migration of the cation in four different states: in addition to probing the influence of the D69^2.50^ protonation state, we examined the role of the conformation of the Y440^7.53^ sidechain, which substantially affects the width and overall shape of the formed hydrated pathway into the cytoplasm (Fig 3C,D).

When D69^2.50^ is charged (Fig 4), the free energy difference between the internal the Na^+^ ion binding site and the free intracellular bulk solution is ~30 kJ/mol. Accordingly, the active conformation of m2r retains a Na^+^ ion at the allosteric site with relatively high affinity, as long as D69^2.50^ remains unprotonated. The major barrier to migration into the cytoplasm is located near the Y440^7.53^ sidechain. In its upward state, the free energy barrier amounts to ~41 kJ/mol, while it increases to ~48 kJ/mol in the downward state (Fig 4).

**Figure 4:**
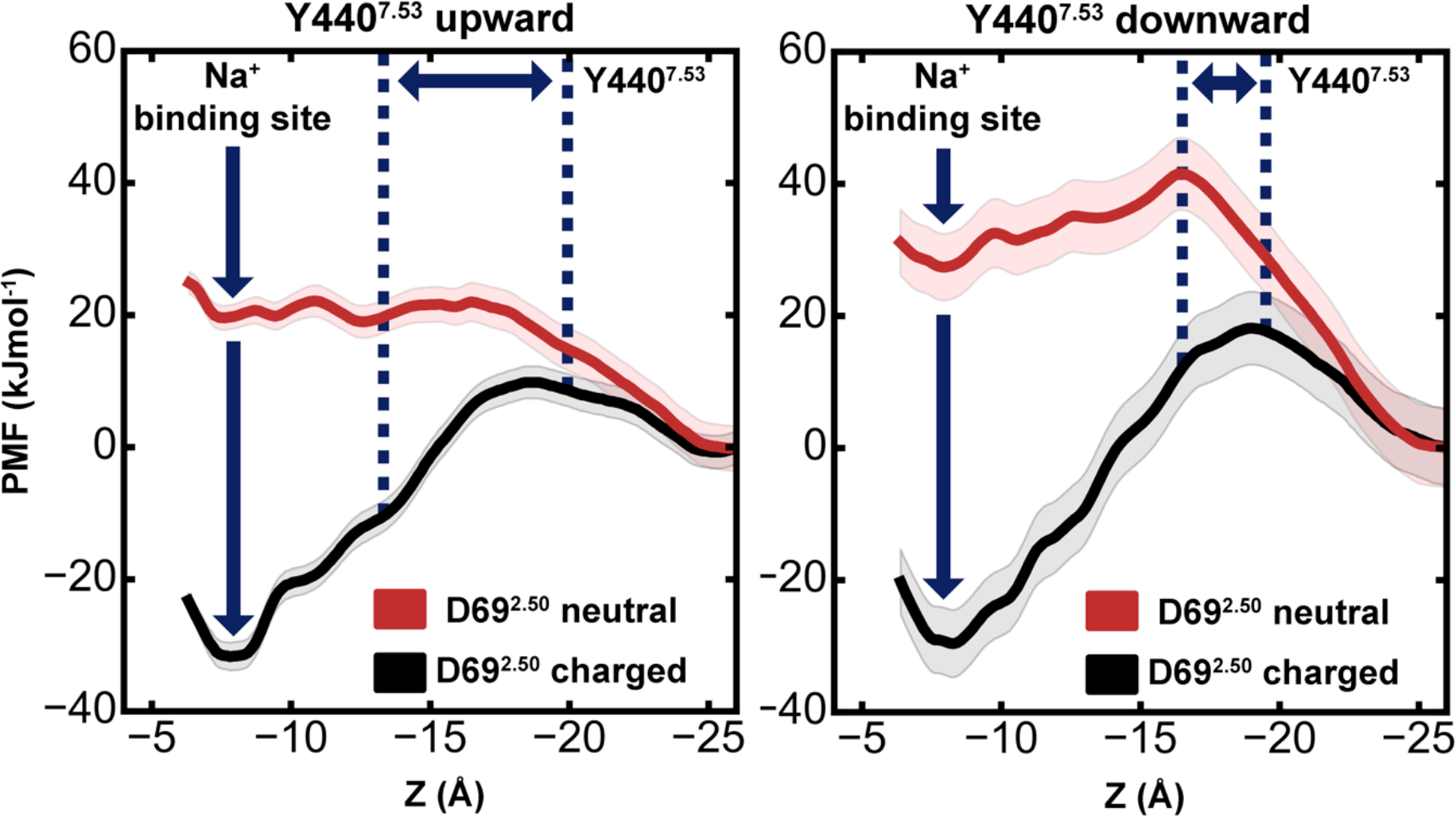
Energetics of Na^+^ translocation from the hydrophilic pocket to the intracellular side. Equilibrium potential of mean force (PMF) profiles of the energetics of Na^+^ translocation along the Z-axis in m2r without any applied voltage or concentration gradients. Four relevant states were considered: **(Left)** negatively charged D69^2.50^ (black) or neutral D69^2.50^ (red) with the Y440^7.53^sidechain in an upward conformation; **(Right)** negatively charged D69^2.50^ (black) or neutral D69^2.50^ (red) with a downward-oriented Y440^7.53^ sidechain. The standard deviation of the PMF, obtained from Bayesian bootstrap analysis, is depicted as shaded area. For each PMF, the intracellular bulk solution was used as a reference, and the range of positions adopted by the Y440^7.53^ sidechain is denoted by blue dotted lines.

As our p*K*_a_ calculations showed that even a moderate displacement of the Na^+^ ion away from its binding site at D69^2.50^ is likely to lead to a protonation change of the aspartate, we also calculated the PMF of the Na^+^ ion movement along the intracellular pathway for neutral D69^2.50^. Importantly, this state no longer shows any affinity for the Na^+^ ion, and ion movement into the intracellular bulk is not obstructed by any energy barrier significantly larger than the thermal energy (*kT*, ~2.5 kJ/mol) in the upward-oriented Y440^7.53^ conformation. When Y440^7.53^ is oriented downward, a small but readily surmountable energy barrier (on physiologically relevant timescales) of ~14 kJ/mol exists for this transition. The downward conformation of Y440^7.53^ in conjunction with the neutral state of D69^2.50^ also has a small influence on the shape and configuration of the ion binding site at D69^2.50^, which leads to a reduction of the number of hydrogen bonds formed between the protein and the ion (Fig S5), raising the free energy of binding at this site further by ~7.5 kJ/mol (Fig 4A, B).

### Conservation of the pocket and intracellular exit channel

Additional support for an important role of intracellular Na^+^ egress in the activation of class A GPCRs is provided by analysis of residue conservation along its exit pathway. As we detailed previously (Katritch et al., 2014), there is a remarkable level of conservation for the 16 residues of the Na^+^ binding pocket in class A GPCRs (Figure 5, Table S1), suggesting conserved functional role of Na^+^ in receptor activation mechanism. Interestingly, our analysis of Na^+^ contacts along the MD trajectories in this study shows that the residues lining the ion exit path to the intracellular side are well conserved too. Thus, out of the 36 contact residues, 32 are 100% conserved among all five muscarinic receptors, 17 are >90% conserved among all aminergic receptors, and 22 are consensus residues among all class A GPCRs. Most importantly, the predicted exit pathway includes Na^+^ contacts with the highly conserved N^1.50^ (100% and 98% conserved in aminergic and in all class A respectively), D^3.49^ (100% and 64%), Y5.58 (94% and 73%) and other residue positions generally conserved as polar residues, including N^1.60^, T^2.37^ and N^2.39^. Particularly, in the inactive M2 muscarinic receptor and in other inactive state GPCR structures as well, the Y^7.53^ residue is directed towards the Na^+^ ion-binding pocket, and hence may play a role as first point of polar contact outside the Na^+^ ion-binding pocket for the intracellular movement of Na^+^. The Na^+^ ion passage towards the cytosol may be further facilitated by other conserved polar residues, including D^3.49^, N^2.39^, N^2.40^ and T^2.37^. Such conservation of the Na^+^ ion pocket and the path for intracellular egress of Na^+^ suggests that Na^+^ transfer described in this study can occur in all muscarinic receptors and other class A GPCRs, comprising a key “irreversible” part of the activation mechanism.

**Figure 5:**
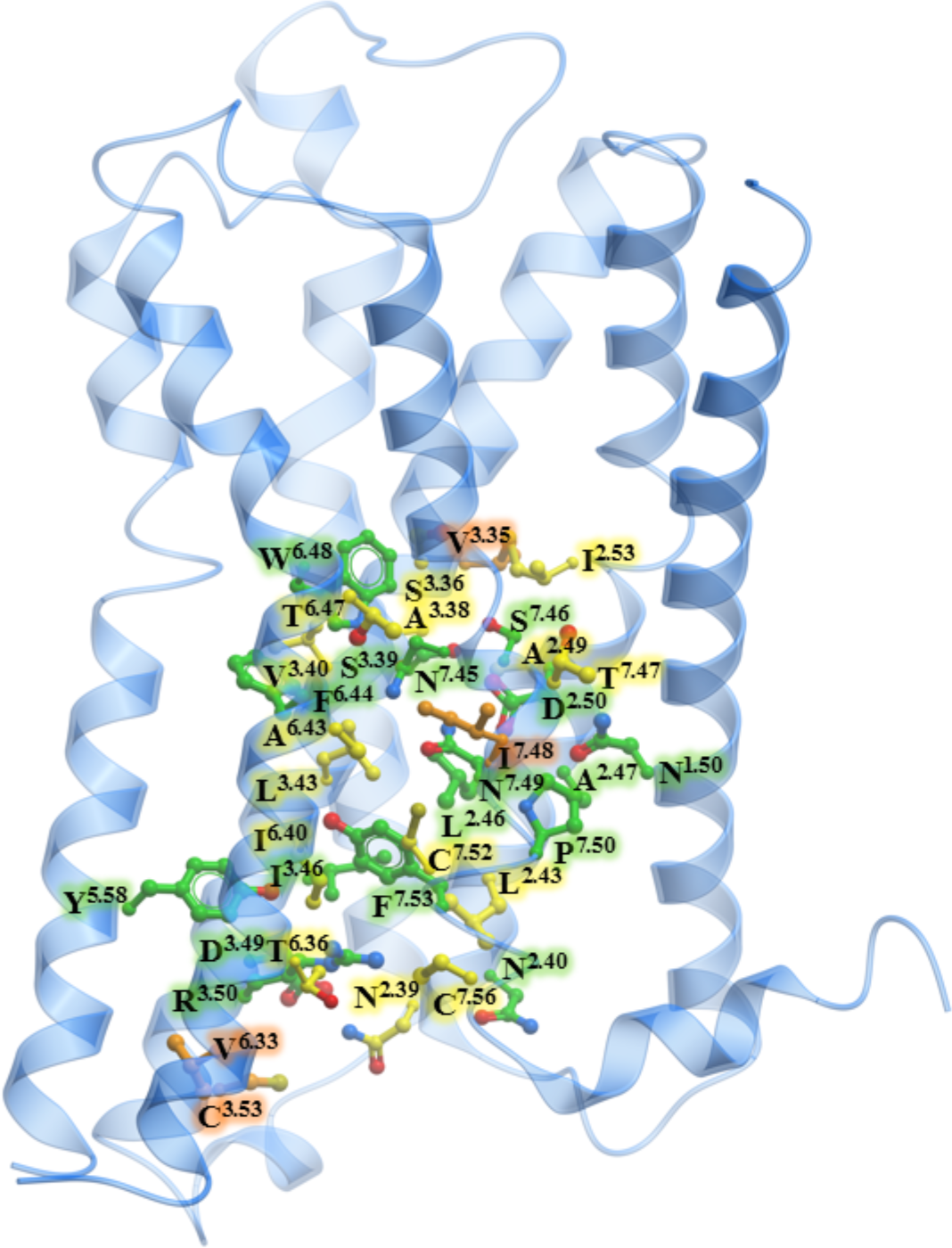
Conservation of the intracellular Na^+^ ion pathway. Muscarinic M2 receptor shown in blue cartoon representation, along with ball-and-stick representation of residues involved in the egress of the Na^+^ ion. The carbon atoms of 17 residues that are >90% conserved among aminergic receptors are shown in green, the carbon atoms of additional 15 residues that are conserved among the muscarinic family of receptors are shown in yellow, the carbon atoms of the 4 non-conserved residues are shown in orange.

## DISCUSSION

The principal role of GPCRs is to transmit information about an extracellular agonist binding event towards the cytoplasm, by catalysing GDP release from a bound intracellular G-protein complex (Pierce et al., 2002). This is known to involve conformational changes in the receptor, including conserved residue microswitches, and large scale movement of TM helices 6 and 7 in the intracellular side that open the nucleotide binding site of the Gα protein (Dror et al., 2015; Huang et al., 2015; Mahoney & Sunahara, 2016). It has, furthermore, long been recognised that G-protein binding, and stabilisation of this conformation on the intracellular side of the receptor, increases agonist affinity on the extracellular face (DeVree et al., 2016; Maguire, Van Arsdale, & Gilman, 1976).

Na^+^ ions, binding to an internal receptor site between the G-protein and the external ligand binding pockets, are known to act as powerful allosteric modulators of class A GPCR function (Katritch et al., 2014; Pert & Synder, 1974). Na^+^ was found to selectively diminish the affinity of agonists, but not antagonists, to GPCR, which can be interpreted as a structural stabilisation of the inactive receptor state by the ions (Miller-Gallacher et al., 2014; Quitterer, AbdAlla, Jarnagin, & Müller-Esterl, 1996; Selley, Cao, Liu, & Childers, 2000). Accordingly, while receptor X-ray structures of sufficient resolution crystallised in the inactive state display a Na^+^ ion bound to D^2.50^, this binding site is collapsed in active receptor conformations, and ions are not observed (Huang et al., 2015; Katritch et al., 2014). Mutations around the Na^+^ ion binding site have a major impact on receptor function in most class A GPCRs either completely abolishing G-protein activation, or resulting in constitutive ligand independent or pathway biased signaling (Fenalti et al., 2014; Liu et al., 2012; Massink et al., 2015).

Our work shows that the Na^+^ ion binding pocket, which is accessible only from the extracellular face in the inactive state (Selent, Sanz, Pastor, & de Fabritiis, 2010; Vickery, Machtens, Tamburrino, Seeliger, & Zachariae, 2016), is transformed into a fully permeable, water-filled channel in the activated receptor conformation of m2r. This channel bridges the extracellular ligand and intracellular G-protein binding sites. Water access from the ligand binding site all the way to the cytoplasmic side of the receptor has previously also been observed in simulations on the A_2A_R and 5-HT_1A_ receptors (Yuan et al., 2014, 2016). We show here that the activated receptor state permits the Na^+^ ion to cross the receptor towards the cytoplasmic side, without experiencing any major energy barriers. The high hydration level of this pathway in the active state is thereby an important factor in facilitating ion passage. The correlation between hydration level and ion transfer has previously been demonstrated in the case of ion channels (Beckstein et al., 2003; Dong, Fiorin, Carnevale, Treptow, & Klein, 2013; Zhu & Hummer, 2012). In simulations of the inactive state, by contrast, the application of substantially larger forces seems to be necessary to achieve inward migration of Na^+^, as no continuous hydrated channel is formed (Shang et al., 2014).

The inward motion of the Na^+^ ion is facilitated by a protonation change of D^2.50^ from the negatively charged to the neutral form, which we show to occur even upon small displacements of the ion from its equilibrium binding position. Neutralisation of D^2.50^ substantially reduces its affinity for Na^+^ ions. Migration of the ion towards the cytosol is then driven by the negative membrane voltage and by a greater than 10-fold Na^+^ gradient across the cytoplasmic membrane under physiological conditions, both strongly attracting Na^+^ ions inwards. Indeed, we observe that moderately negative membrane voltages allow fast escape of the allosteric Na^+^ ion to the cytoplasm on 10–100 ns-timescales in our simulations.

According to our results, conformational changes associated with agonist binding from the extracellular side and/or G-protein binding from the cytoplasm alter the Na^+^ site conformation and the dynamics of the Na^+^–D^2.50^ salt bridge. This, in turn, leads to a protonation change of this residue, and subsequent egress of the Na^+^ ion via a hydrated exit channel to the intracellular side. We therefore suggest that intracellular Na^+^ ion transfer, facilitated by the membrane potential and Na^+^ gradient, is a pivotal step during receptor activation, as it traps the receptor in the active state (Fig 6). It has been shown that, once activated, GPCRs remain in a prolonged active state, capable of signalling even when the receptors are internalized from the cytoplasmic membrane during endocytosis (Irannejad et al., 2013; Thomsen et al., 2016). The crucial role of the Na^+^ ion movement within the receptor is reflected by the nearly complete conservation of the Na^+^ ion binding site in class A GPCRs as well as the high conservation level of the exit pathway. The mechanism suggested here is also consistent with agonist independent basal signalling of GPCRs (Kobilka & Deupi, 2007), explaining this phenomenon as spontaneous protonation of D^2.50^ and egress of the bound Na^+^ ion on the intracellular pathway, leading to receptor activation.

**Figure 6:**
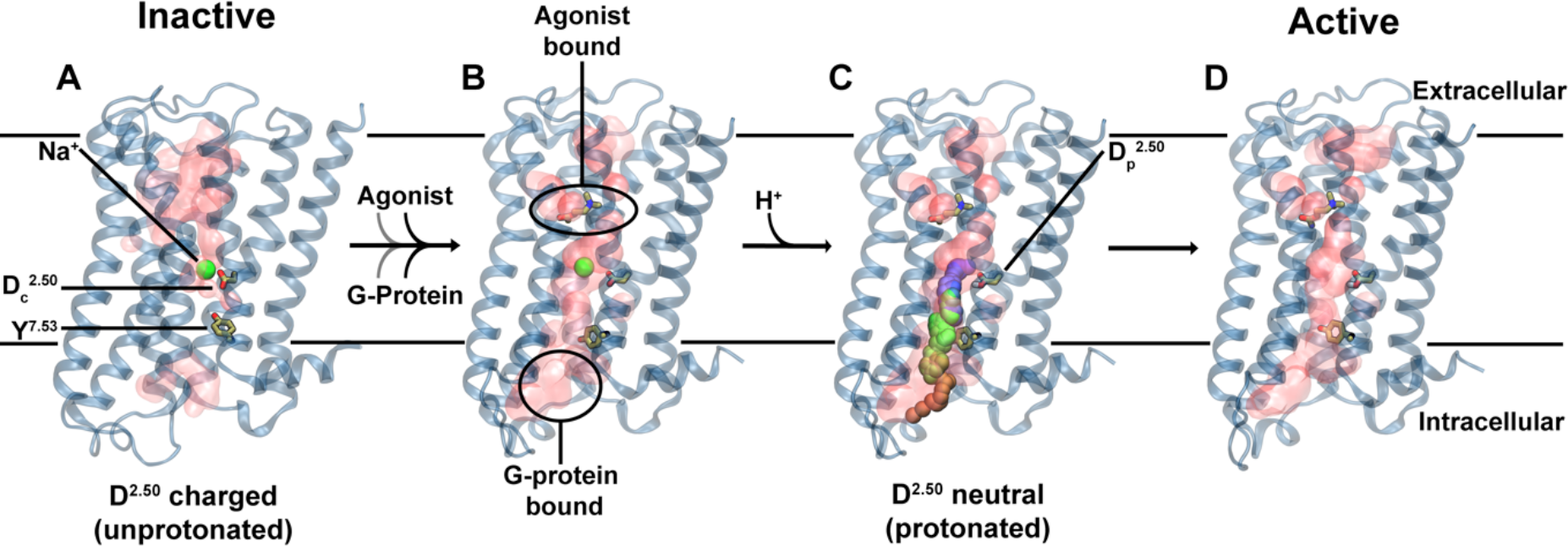
Proposed role of Na^+^ translocation in GPCR activation. Key checkpoints during the transition from the inactive (A) to active (D) state of the receptor. **(A)** The initial, inactive receptor conformation shows no bound agonist or G-protein, and displays a Na^+^ ion bound in a pocket which is sealed towards the cytosol by a hydrophobic layer around Y^7.53^. **(B)** G-protein and agonist bind to the receptor (in undetermined order), leading to the formation of a continuous water channel across the GPCR. The increased mobility of the Na^+^ ion results in a pK_a_ shift and subsequent protonation of D^2.50^. **(C)** Neutralisation of D^2.50^ and the presence of the hydrated pathway facilitate transfer of Na^+^ to the intracellular side, which is driven by the transmembrane Na^+^ gradient and the membrane voltage. **(D)** The expulsion of Na^+^ towards the cytosol results in a prolonged active state of the receptor.

Charge movements within membrane proteins, such as the coupled transfer of Na^+^ ions and protons suggested by our MD simulations and p*K*_a_ calculations, should be sensitive to the membrane voltage. Indeed, it has been demonstrated that GPCR signalling is modulated by membrane voltage changes (Ben-Chaim et al., 2006; Mahaut-Smith, Martinez-Pinna, & Gurung, 2008; Martinez-Pinna, Tolhurst, Gurung, Vandenberg, & Mahaut-Smith, 2004; Moreno-Galindo, Alamilla, Sanchez-Chapula, Tristani-Firouzi, & Navarro-Polanco, 2016; Rinne et al., 2015; Vickery, Machtens, Tamburrino, et al., 2016). This applies both to the conformation of the receptors as well as their transmitted signal. Our findings are therefore consistent with these observations, as they suggest that movement of ions in the receptors constitute a key element in the receptor activation process. The observed voltage regulation of GPCRs is of particular relevance for receptors expressed in electrically excitable cells (Heifetz, James, Morao, Bodkin, & Biggin, 2016). In these cell types, the membrane voltage undergoes large-scale oscillations during action potentials. The transmitted receptor signal could thereby be tuned depending on the specific cell type and its excitation status (Vickery, Machtens, & Zachariae, 2016). Crucially, many GPCR drug targets are located in excitable tissue in the brain or muscle, where voltage regulation and a differential response to drugs may play an important role.

To summarise, our results suggest a model for class A GPCR activation, in which conformational changes induced by G-protein and agonist binding are accompanied by the intracellular transfer of an internally bound Na^+^ ion. Importantly, these conformational changes encompass rearrangement of the sidechain of Y^7.53^, a conserved receptor microswitch (Katritch et al., 2013), which in its upward state allows nearly barrier-free intracellular permeation of Na^+^ ions. This observation forms a functional link between the major Na^+^ binding site D^2.50^ and Y^7.53^ as the first polar point of contact on the intracellular migration pathway of the Na^+^ ion. Translocation of the ion is further facilitated by protonation of the conserved D^2.50^ residue (Fig 6), and driven by the physiological membrane Na^+^ and voltage gradients. The voltage sensitivity of GPCRs, which has been previously reported for many receptors (Vickery, Machtens, & Zachariae, 2016), would thus be a natural consequence of an activation mechanism which incorporates the movement of ions as a key element. The Na^+^ free receptors are likely to be trapped in an active state, potentially explaining the prolonged mechanisms of signalling observed in many GPCRs.

## METHODS

The simulation systems for the m2r were constructed using the crystal structures (PDB: 3UON, 4MQT) (Haga et al., 2012; Kruse et al., 2013). Ligands and non-GPCR subunits were removed. The missing loop ICL3 was modelled using Modeller (v9.14)(Šali & Blundell, 1993). For both simulation systems all internal water molecules and ions were retained. The charged N- and C-termini were capped using acetyl and methyl moieties, respectively. All ionisable groups were simulated with default protonation states, unless otherwise mentioned. Each receptor was embedded into an equilibrated and hydrated 1,2- palmitoyl-oleoyl-sn-glycero-3-phosphocholine (POPC) lipid bilayer using the GROMACS utility g_membed (Wolf, Hoefling, Aponte-Santamaría, Grubmüller, & Groenhof, 2010) resulting in a system size of ~92 x 88 x 97 Å. A concentration of 150 mM NaCl in the aqueous solution was used for the single bilayer systems. During equilibration, all protein heavy atoms were position-restrained with a force constant of 1000 kJ mol^-1^ nm^-2^ for 5-10 ns. Due to the low degree of internal hydration and medium resolution of the m2r structures the equilibrations were extended by another 100 ns, now without position restraints, to enable full hydration of the hydrophilic pocket. To study the active structure, the ligand carbachol was parameterised using AMBER16, GAFF2, AM1-BCC parameters (Case et al., 2016), and docked into the orthosteric ligand binding site using GOLD (v5.2.2). A Na^+^ ion was placed into the hydrophilic pocket in the inactive structure. We used a targeted MD approach with the RMSD to the protein backbone of the active m2r crystal structure (PDB: 4MQT) as a reference, in order to gently enforce the transition from the inactive (PDB: 3UON) to the active state within ~250 ns. The two major conformations of Y440^7.53^ we observed during this simulation were probed systematically in the PMF calculations using distance restraints between N^1.50^-C_α_ and D^2.50^-C_α_ to Y^7.53^-C_**ζ**_ or dihedral restraints on the sidechain of Y^7.53^. To keep the G-protein binding site in an active conformation despite the absence of bound G-protein, we applied a minimal set of four distance restraints to the C_α_ atoms of the terminal groups of TM helices 2,5,6 and 7, namely residues 2.39-6.33, 2.39-5.61, 2.43-7.54 and 6.36-7.54, at this interaction site (Fig S6).

For the CompEL simulations, the aforementioned active systems were duplicated along the z axis to construct double bilayer systems. A NaCl gradient of 150mM:10mM between the extracellular and intracellular compartments was used, along with an ion imbalance of 1 to 2 Cl^-^ ions to generate a V_m_ of ~-250 to ~ -500mV, as previously described (Kutzner, Grubmüller, de Groot, & Zachariae, 2011). The V_m_ was determined by the GROMACS utility gmx potential.

To calculate the PMF for the Na^+^ ion within the hydrophilic pocket at neutral V_m_, umbrella sampling calculations were performed in bins of 0.25Å and analysed with the GROMACS utility gmx wham. We used a simulation time of 50ns in each window and harmonic potentials of 900–2000 kJ mol^-1^ nm^-2^ to restrain the Na^+^ ion in the z-direction. The standard deviation of the PMF profiles was estimated by using the Bayesian bootstrap method, as implemented in gmx wham, with 200 runs. The free energy of the Na^+^ ion in bulk solution was set to 0. The position of the Na^+^ ion (Z-coordinate) is reported relative to the D103^3.32^- C_α_ atom (ligand binding site).

For all simulations, the amber99sb_ildn force field was used for the protein (Lindorff-Larsen et al., 2010), Berger parameters for lipids (Berger, Edholm, & Jähnig, 1997), which were adapted for use with the amber99sb force field (Cordomí, Caltabiano, & Pardo, 2012), and the SPC/E model for water molecules (Berendsen, Grigera, & Straatsma, 1987). Water bond angles and distances were constrained by SETTLE (Miyamoto & Kollman, 1992) while all other bonds were constrained using the LINCS method (Hess, Bekker, Berendsen, & Fraaije, 1997). The temperature and pressure were kept constant throughout the simulations at 310 K and 1 bar, respectively, with the protein, lipids, and water/ions coupled individually to a temperature bath by the v-rescale method using a time constant of 0.2 ps and a semi-isotropic Berendsen barostat (Berendsen, Postma, van Gunsteren, DiNola, & Haak, 1984; Bussi, Donadio, & Parrinello, 2007). Employing a virtual site model for hydrogen atoms (Feenstra, Hess, & Berendsen, 1999) allowed the use of 4-fs time steps during the simulation. All simulations were performed with the GROMACS software package, version 5.1.2

The p*K*_a_ calculations were performed using a continuum electrostatics method, namely the Poisson-Boltzmann/Monte Carlo (PB/MC) approach, on multiple snapshots taken at a 2 ns interval from different umbrella sampling simulations. PB calculations were performed using MEAD (version 2.2.9)(Bashford & Gerwert, 1992) with a dielectric constant (ε_p_) of 4 for the protein and 80 for the solvent (ε_w_), in the presence of an explicit membrane. The temperature was set to 310 K and the ionic strength to 0.145 M. The same temperature was used for MC calculations (10^3^ steps in each calculation), which were performed using MCRP (Baptista, Martel, & Soares, 1999). Each MC step consisted of a cycle of random choices of a state for all individual sites and pairs of sites with couplings above 2.0 p*K*_a_ units (Baptista et al., 1999), whose acceptance/rejection followed a Metropolis criterion (Metropolis, Rosenbluth, Rosenbluth, Teller, & Teller, 1953); tautomeric forms were not included.

The GROMACS software package, version 5.0.4 analysis toolkit was used to identify residues with non-hydrogen heavy atoms within 4 Å of the sodium ion path during the simulations. The residue conservation profile of the amino acids was obtained from the GPCRdb server (Isberg et al., 2015).

## ACKNOWLEDGEMENTS

This work was supported by the BBSRC (Training Grant BB/J013072/1 to U.Z.) and the Scottish Universities’ Physics Alliance (C.A.C, A.V.P. and U.Z.). This research was partially supported by National Institute of Health grant DA035764 to V.K. We thank Salomé Llabrés and Daniel Seeliger for fruitful discussions.

## SUPPLEMENTAL INFORMATION

Figure S1: Backbone RMSD during a targeted MD simulation from the inactive to the active state of m2r.

Figure S2: Na^+^ position during a targeted MD simulation from the inactive to the active state of m2r.

Figure S3: Y440^7.53^ conformations used in p*K*_a_ and PMF calculations.

Figure S4: Backbone RMSD during MD simulations of the active state m2r under membrane voltage.

Figure S5: Number of hydrogen bonds around the Na^+^ binding site.

Figure S6: Depiction of the minimal set of distance restraints used to maintain the active conformation of the m2r at the G-protein binding site.

Figure S7: Na^+^ ion migration across the receptor to the intracellular side.

Table S1. Conservation of the residues in the transmembrane region that were observed to be in contact (>4.5 Å) with Na^+^ in the MD simulations. Overall, the sodium ion was observed to come into close proximity with 34 residues.

